# anndataR improves interoperability between R and Python in single-cell transcriptomics

**DOI:** 10.1101/2025.08.18.669052

**Authors:** Louise Deconinck, Luke Zappia, Robrecht Cannoodt, Martin Morgan, scverse core, Isaac Virshup, Chananchida Sang-Aram, Danila Bredikhin, Brian Schilder, Ruth Seurinck, Yvan Saeys

## Abstract

**Summary:** Many single-cell transcriptomics datasets are stored in the HDF5-backed AnnData (H5AD) file format, as popularised by the Python scverse ecosystem. However, accessing these datasets from R, allowing users to take advantage of the strengths of each language, can be difficult. anndataR facilitates this access by allowing users to natively read and write H5AD files in R, convert them to and from SingleCellExperiment or Seurat objects, or even work with the resulting R AnnData object directly. We perform rigorous testing to ensure compatibility between Python-written and R-written H5AD files, guaranteeing long-term interoperability between languages.

**Availability:** anndataR’s source code is available on GitHub at scverse/anndataR under the MIT license. It is compatible with R version 4.5, has been archived at 10.5281/zenodo.18775712 and included within Bioconductor: 10.18129/B9.bioc.anndataR. Installation instructions and tutorials can be found in the online documentation at anndatar.scverse.org. Issues can be reported at the GitHub repository. Code to reproduce the analyses performed can be found on GitHub at LouiseDck/anndataR-paper, archived at 10.5281/zenodo.18792241.

## 1. Introduction

In the single-cell transcriptomics field, three main analysis ecosystems exist: scverse (Virshup et al., 2023), Seurat (Satija et al., 2015) and Bioconductor (Amezquita et al., 2020), each defining its own in-memory data format. Two of these, the SingleCellExperiment object used by Bioconductor and the Seurat object, are implemented in R, while the scverse AnnData object (Virshup et al., 2024) is Python based.

Each ecosystem has specific advantages for single-cell transcriptomics (scRNA-seq) data analysis and best practices and benchmarks recommend combining tools from different ecosystems to obtain an optimal processing pipeline for scRNA-seq data (Heumos et al., 2023). For instance, Seurat easily accommodates multi-modal assays, while Bioconductor comes with easy access to extensive statistical tooling and scverse leverages scalability and access to machine learning ecosystems. On top of that, small implementation differences in basic functionality (e.g. PCA) between the ecosystems may produce significantly different results (Rich et al., 2024). As a result, users may need to switch between data formats or even programming languages to perform different analysis steps or to replicate existing analyses. Unfortunately, this is not straightforward and a number of issues arise when trying to convert data between formats. These issues stem from a) structural differences between the data formats and b) the different programming languages.

### 1.1. Differences in structure

All three data formats were developed to work with single-cell transcriptomics data but they take different approaches and structure their objects differently. An overview of the structure of these objects can be found in Table 1.

**Table 1.** Overview of the three different data formats and the slots where specific information is stored.

| Stored information | AnnData | SCE | Seurat v5 |
| --- | --- | --- | --- |
| Expression matrices | X, layers | assays | layers |
| Unfiltered expression matrices (different features) | raw | altExp | layers |
| Paired measurements (spike in, CITE-seq, ...) |  | altExp | assays |
| Multi-omics data (different cells) |  |  | assays |
| Cell-level metadata (cell type, ...) | obs | colData | cell level metadata |
| Feature-level metadata (number of cells, ...) | var | rowData | assay level metadata |
| Multidimensional cell-level metadata (dimensionality reductions) | obsm | reduced-Dims | reductions |
| Multidimensional feature-level metadata (loadings of dimensionality reductions) | varm | reduced-Dims | reductions |
| Cell by cell similarity matrices | obsp | colPairs | graphs |
| Feature by feature similarity matrices | varp | rowPairs |  |
| Other unstructured metadata | uns | metadata | misc |

For example, the AnnData object contains a `varm` slot which stores multidimensional variable annotations (such as PCA loadings) but SingleCellExperiment and Seurat instead store this information as metadata of the cell reduction objects. The Seurat object also has no corresponding slot for the AnnData varp slot, which is used for pairwise variable level annotations. Seurat and SingleCellExperiment can also store multimodal data, which is not possible in AnnData. Additionally, the frameworks differ in matrix orientation conventions. AnnData uses an observations by features layout, with observations (cells) in rows and features (genes) in columns. Conversely, SingleCellExperiment and Seurat implement these matrices with cells as columns and genes as rows. Because of these structural differences, in-depth knowledge of each object and the related storage conventions is required to correctly convert between formats, which presents a barrier to users moving between ecosystems.

### 1.2. Different programming languages

The different programming languages also complicate interoperability. Seurat and Bioconductor are written in R while scverse is written in Python. As mentioned earlier, tools in both programming languages are required to optimally analyse a single dataset. To address this, two other core data structures within the scverse ecosystem have some cross-language interoperability: MuData (Bredikhin et al., 2022), designed for multimodal data, provides R and Julia implementations, and SpatialData (Marconato et al., 2025) maintains active collaboration with R developers (Marconato et al., 2024). For single-cell transcriptomics analysis, there are many software packages that facilitate the conversion between different objects (Table 2). They use a combination of approaches to transfer data between programming languages: they either use a so-called foreign function interface (FFI) to transfer information between languages during an interactive session or they write and read the data to and from disk.

**Table 2.** Overview of packages aiming to provide some form of interoperability between AnnData, SingleCellExperiment and Seurat objects.

| Tool | Prog.<br>language | FFI | File<br>format | Conversion |
| --- | --- | --- | --- | --- |
| dynverse/<br>anndata<br>(Cannoodt,<br>2025) | R | ✓ |  |  |
| anndata2ri<br>(Angerer et al.,<br>2020) | Python | ✓ |  | AnnData↔SCE |
| Loom<br>(Linnarsson<br>Lab, 2024;<br>Hoffman and<br>Satija, 2018) | R, Python |  | Loom<br>(h5) |  |
| scDIOR (Feng<br>et al., 2022) | R, Python |  | scDIOR<br>(h5) | AnnData↔SCE<br>AnnData↔Seurat |
| BiocPy with<br>rds2py<br>(Kancherla,<br>2025) | Python |  |  |  |
| alabaster<br>dolomite (Lun,<br>2024; Lun and<br>Kancherla,<br>2025) | R, Python |  | HDF5,<br>csv |  |
| SeuratDisk<br>(Hoffman,<br>2023) | R |  | h5Seurat | Seurat↔AnnData |
| zellkonverter<br>(Zappia and<br>Lun, 2025) | R | ✓ |  | AnnData↔SCE |
| sceasy (Cakir<br>et al., 2020) | R | ✓ |  | Converts between<br>AnnData, SCE,<br>Seurat and Loom |
| schard (Mazin,<br>2025) | R |  |  | Converts between<br>AnnData, SCE and<br>Seurat<sup>1</sup> |
| anndataR (this<br>paper) | R |  |  | Converts between<br>AnnData, SCE and<br>Seurat |
| SCUBA<br>(Showers et al.,<br>2025) | R | ✓ | provides<br>API |  |
| scKirby<br>(Schilder, 2023) | R | ✓ |  | Converts between<br>AnnData, SCE,<br>Seurat and Loom<sup>2</sup> |

FFIs like reticulate (Ushey et al., 2025) or rpy2 (Gautier, 2025) allow a program written in one language to call or use functions written in another language. This allows users to flexibly call foreign functions, transferring data from the native language to the foreign language as needed. However, such an approach faces several limitations. It is limited to built-in datatypes (such as vectors, lists or integers), requires managing both a Python and R environment, and naive use of these interfaces can lead to excessive memory usage. Software packages that use FFI to facilitate interaction with single-cell specific data formats include anndata2ri (Angerer et al., 2020) and dynverse/anndata (Cannoodt, 2025).

The second approach, working on an object that gets intermediately stored on disk, requires a standard on-disk format accessible from both languages. A shared standard, loom, was introduced but was not fully taken up by the community. The AnnData HDF5-based (The HDF Group) H5AD format has become the de facto standard for Python users and in some public repositories while R objects are commonly saved as non-interoperable binary Rds files. To address this, packages such as rds2py (Kancherla, 2025), alabaster (Lun, 2024) and dolomite (Lun and Kancherla, 2025) provide ways to read and write SingleCellExperiment objects in Python. SeuratDisk (Hoffman, 2023) provides a way to save Seurat objects as interoperable HDF5 files.

Other packages, including sceasy (Cakir et al., 2020) and zellkonverter (Zappia and Lun, 2025), combine these approaches to make H5AD files useable in R. They allow reading and writing of H5AD files to and from Seurat or SingleCellExperiment object but do so by using the reticulate software package, an FFI, to read the H5AD file using the Python anndata package before converting formats. This approach only partially alleviates the burden of working with these FFIs. R users (or package maintainers) still need to maintain a Python environment and be mindful of in-memory duplication of data, even when all their analysis is performed in R. The data type conversions from Python to R can also be more cumbersome than those needed when reading in the HDF5 file from disk.

## 2. Results

To overcome these limitations we introduce anndataR, a new package which allows native reading and writing of H5AD files in R, without requiring a Python environment, provides an R AnnData object and facilitates conversion to and from other data formats. anndataR was conceived and developed as a collaborative effort between members of both the scverse and Bioconductor communities, who together identified the gaps in the current interoperability climate.

anndataR is unique in its ability to read and write H5AD files natively in R and convert them to SingleCellExperiment or Seurat objects. By avoiding FFIs, anndataR avoids many of the associated challenges. The conversion functionality it offers bridges the gap between the ecosystems, with sensible defaults (see Supplementary Figure 2 and 3) and fine-grained control for advanced use. anndataR allows users direct access to an R AnnData representation, making it possible to interact with the object directly, in order to solve conversion issues and to facilitate easy extraction of relevant parts of the data.

## 3. Software Design

Users primarily interact with anndataR by reading in an existing H5AD file and specifying whether to return an AnnData object in R (either in memory or HDF5-backed), or immediately perform a conversion to a SingleCellExperiment or Seurat object, as shown in Supplementary Figure 1. The resulting R object can then be used as normal and can be saved to a new H5AD file on completion of the analysis. An example of how anndataR can be used in an analysis workflow is shown in Figure 1.

**Fig. 1:**
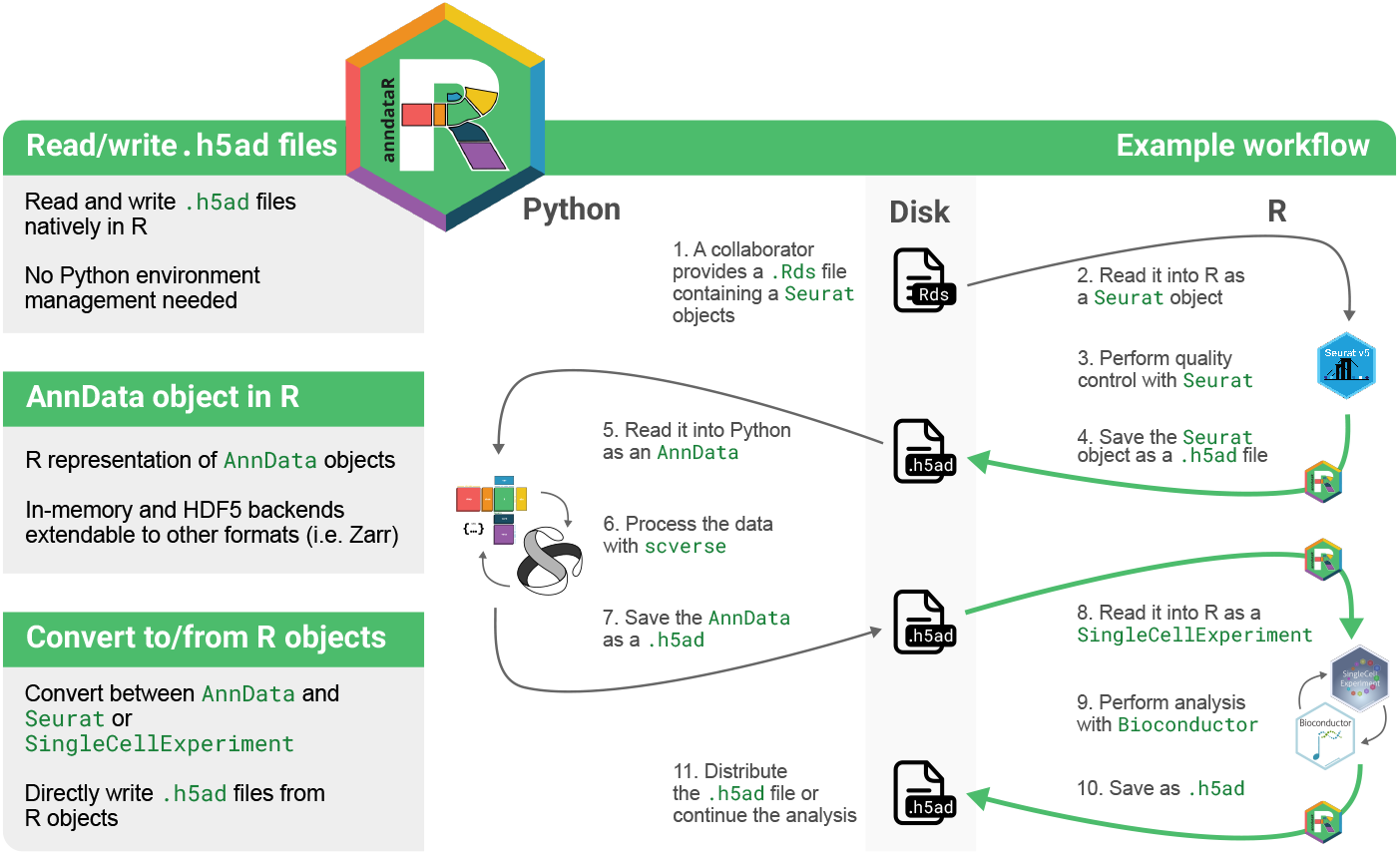
Feature overview and showcase of how to use anndataR in an analysis workflow.

To enable this workflow, we use rhdf5 (Fischer et al., 2025), an R interface to the HDF5 library which handles the low-level interactions with HDF5 files to natively read and write the H5AD format according to the AnnData on-disk specification. To take advantage of existing R analysis packages users also need to be able to convert between AnnData objects and SingleCellExperiment or Seurat objects. By default the anndataR conversion functions try to convert as much data as possible, but also allow the experienced user to completely specify the details of the conversion by providing a mapping for each slot in the resulting object. Full details of the conversion can be found in the package documentation.

We are aware that small changes in any of these data formats can quickly lead to failures in reading, writing or converting. To ensure the package functionality is robust against data format changes, we perform extensive testing. H5AD files written with anndataR are validated against the ones produced by the Python anndata package. This is necessary because there are differences in how Python and R handle data, such as row-major versus column-major matrices, and in how the Python and R HDF5 libraries translate data to H5 datatypes. We perform these checks using a number of round-trip tests for each AnnData slot, checking whether an R-written H5AD slot can be read in Python and vice versa. We also check for differences in the generated H5AD files using the `h5diff` utility. Additionally, we verify the conversions to and from SingleCellExperiment and Seurat by slot-by-slot verification of the objects. This approach to exhaustively testing functionality has been key for the development of anndataR, allowing us to document known limitations and incompatibilities. This has led to a comprehensive and robust tool, (Supplementary Table 1) that is faster and more memory-efficient than comparable tools, particularly for larger datasets (Supplementary Figure 4). Continuous benchmarking via Bencher (https://bencher.dev/perf/anndatar/plots) further ensures that performance does not regress over time.

Finally, in order to represent the complex AnnData structure and replicate the Python interface, make use of inheritance, keep memory usage low and use reference semantics, we chose to use the R6 object oriented class system.

## 4. Conclusion

anndataR meaningfully adds to the interoperability landscape between R and Python for single-cell transcriptomics by allowing users to work natively with H5AD files in R and providing conversion functionality to and from SingleCellExperiment and Seurat objects. This conversion functionality not only provides reasonable defaults but also allows fine-grained user control. Additionally, the package functionality, such as compatibility between R-written and Python-written H5AD files, is rigorously tested.

Finally, the modular design of the software makes it easy to extend the functionality of the package, enabling future support for additional file formats (e.g., Zarr) and other modalities, such as scATAC-seq or CITE-seq (via MuData (Bredikhin et al., 2022)) and spatial data (via the SpatialData (Marconato et al., 2025) framework).

## 5. Acknowledgments

This work was supported by the Research Foundation - Flanders (FWO) [1SF3822N to L.D.]; Chan Zuckerberg Initiative Essential Open Source Software for Science grant [EOSS6-0000000743 to L.Z. and R.C.]; Chan Zuckerberg Initiative Foundation (CZIF) - CZI Seed Networks [CZIF2019-002443 to M.M.]; Ghent University Special Research Fund [BOF21-DOC-105 to C.S.] and the Flanders AI Research (FAIR) Program [174B09119 to R.S. and Y.S.];

We acknowledge the contribution of the scverse core. Members: Can Ergen^1,2^, Danila Bredikhin^3^, Emma Dann^3^, Giovanni Palla^4^, Gregor Sturm^5^, Ilan Gold^6^, Isaac Virshup^4,6^, Jennifer A. Foltz^7^, Luca Marconato^8,9^, Lukas Heumos^10^, Mikaela Koutrouli^11^, Pau Badia-i-Mompel^3^, Philipp Angerer^12^, Roshan Sharma^13,14^, Sara Jimenez^14^, Severin Dicks^10,15^, Tim Treis^12^, Wouter-Michiel Vierdag^16,17^.

^1^Department of Electrical Engineering and Computer Sciences, University of California Berkeley, Berkeley, CA, US ^2^Department of Medicine, University Medical Center Hamburg-Eppendorf, Hamburg, Germany. ^3^Department of Genetics, Stanford University, Stanford, CA, USA ^4^Chan Zuckerberg Initiative, Redwood City, California, USA ^5^Boehringer Ingelheim International Pharma GmbH & Co KG, 88397 Biberach/Riss, Germany ^6^Institute of Computational Biology, Helmholtz, Center Munich, Munich, Germany ^7^Washington University School of Medicine, St. Louis, Missouri, US ^8^European Molecular Biology Laboratory, Genome Biology Unit, Heidelberg, Germany ^9^Division of Computational Genomics and System Genetics, German Cancer Research Center, Heidelberg, Germany ^10^Helmholtz Zentrum München: Munich, Bavaria, DE ^11^Computational Sciences-Center of Excellence, Genentech, South San Francisco, CA, United States ^12^Helmholtz Zentrum München Deutsches Forschungszentrum für Gesundheit und Umwelt: Neuherberg, Bayern, DE ^13^Computational and Systems Biology Program, Sloan Kettering Institute, Memorial Sloan Kettering Cancer Center, New York, NY ^14^Single-cell Analytics Innovation Lab, Memorial Sloan Kettering Cancer Center, New York, NY ^15^Nvidia, Santa Clara, CA, USA ^16^European Molecular Biology Laboratory, Genome Biology Unit, Heidelberg, Germany ^17^Collaboration for joint PhD degree between EMBL and Heidelberg University, Faculty of Biosciences, Heidelberg, Germany

## Footnotes

1 Incomplete: only reads and converts X, obs, var, obsm

2 Combines conversion functionality from other packages

## Notes

https://github.com/scverse/anndataR/

https://github.com/LouiseDck/anndataR-paper

### Competing Interest Statement

The authors have declared no competing interest.

### Summary of Updates

- Added runtime and memory usage analysis - Added a detailed, slot-by-slot comparison of anndataR's functionality against existing tools - Additional diagrams and context detailing exactly how information is mapped between AnnData, Seurat, and SingleCellExperiment objects

