## Supplementary for "anndataR improves interoperability between R and Python in single-cell transcriptomics"

### Supplementary Figure 1

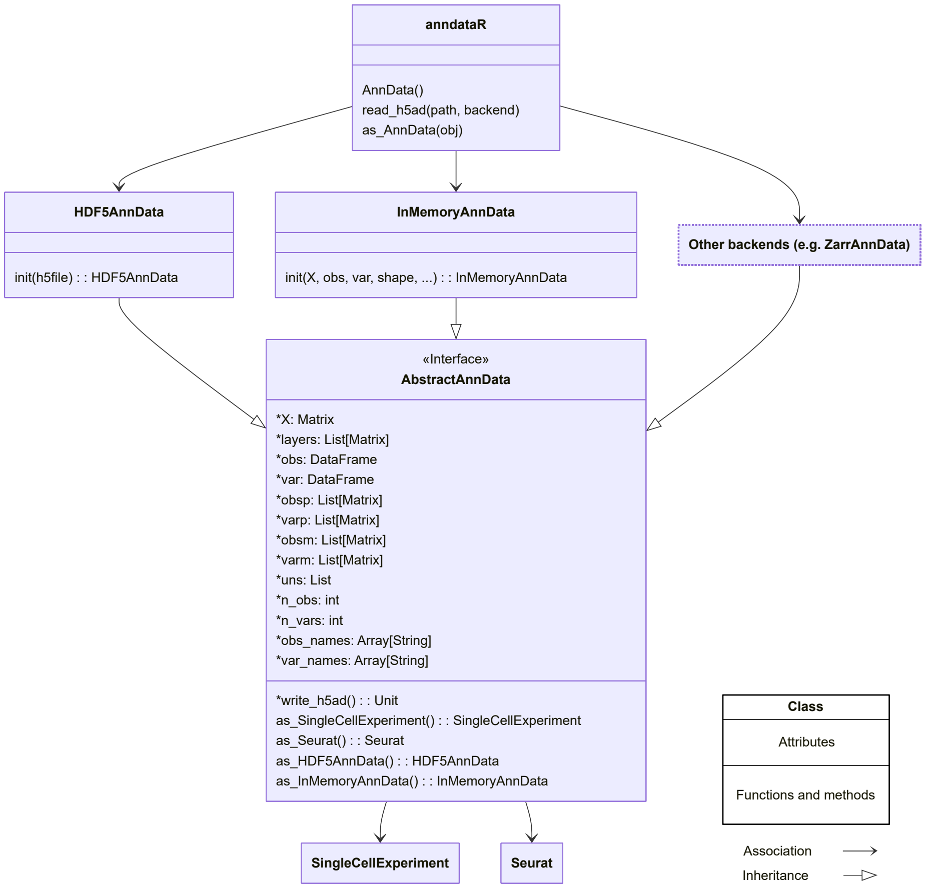


Figure 1: Schematic class diagram of anndataR.

### Supplementary Figure 2

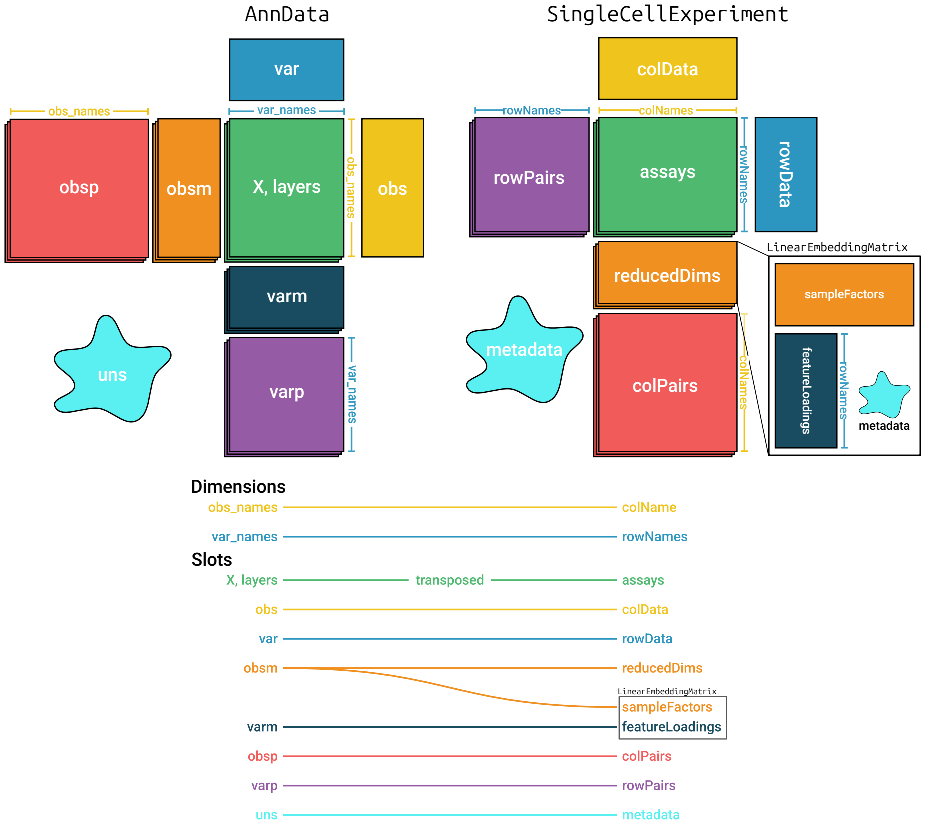


Figure 2: Schematic overview of the default conversion to and from AnnData and SingleCellExperiment objects.

#### Supplementary Figure 3

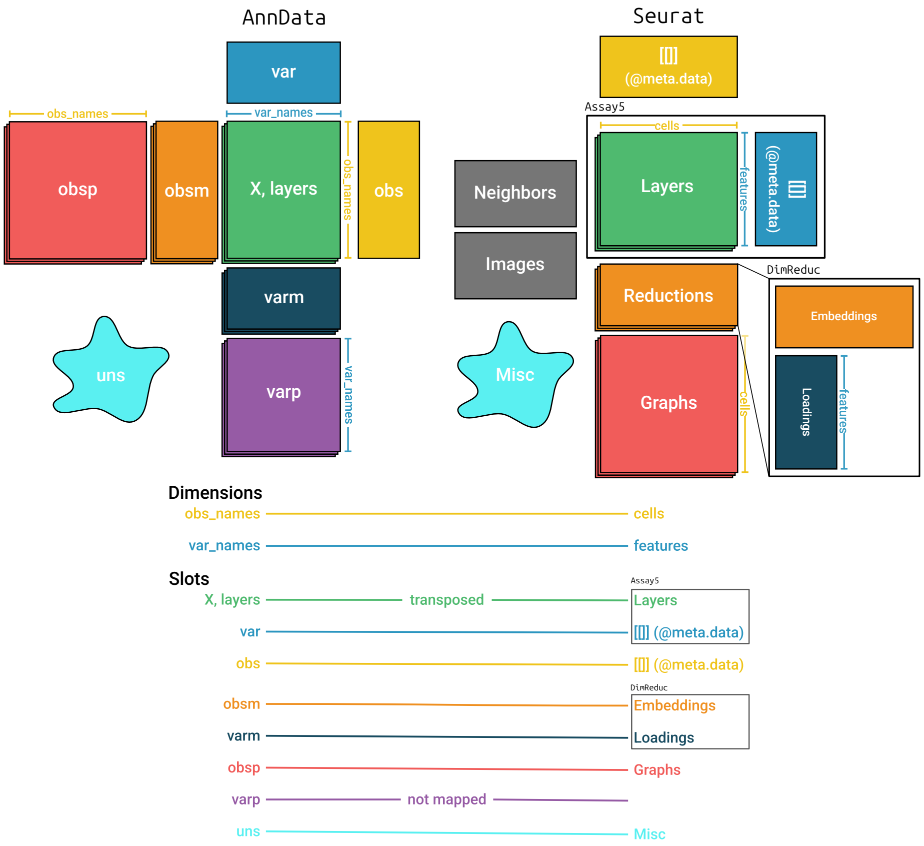


Figure 3: Schematic overview of the default conversion to and from AnnData and Seurat objects. Note that by default, the AnnData `varp` slot will not be converted.

### Supplementary Figure 4

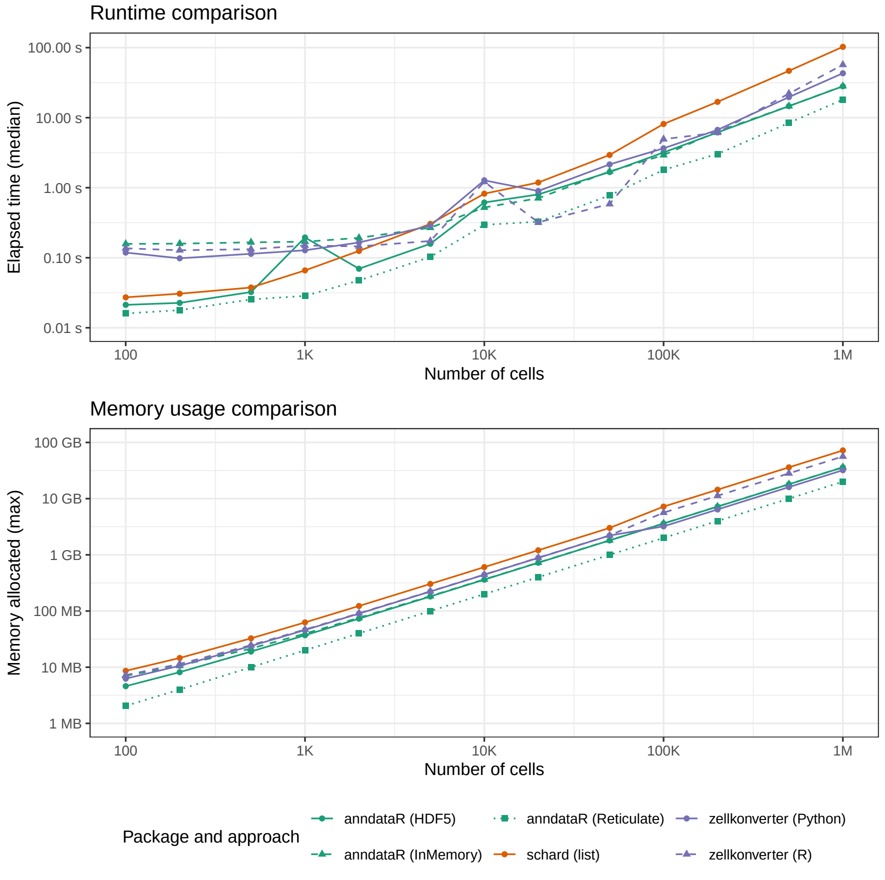


Figure 4: Runtime and memory benchmark for reading an H5AD file and computing the sum of the expression matrix. **Top:** Median elapsed time. **Bottom:** Maximum memory allocated. We compared three anndataR backends (InMemory, HDF5, and Reticulate), two zellkonverter backends (native R reader and Python/basilisk reader), and schard (list export). Synthetic datasets were generated with `dummy_anndata` containing 20 000 genes and between 100 and 1 000 000 cells at approximately logarithmic intervals, with a sparse expression matrix (density 5%). Each benchmark iteration reads the H5AD file and computes the sum of the X matrix; anndataR and schard access the X slot directly, whereas zellkonverter returns a SingleCellExperiment object and is therefore expected to incur slightly higher overhead. Timings report the median of at least 3 iterations. Benchmarks were run on a machine with an AMD Ryzen 9 5950X (16 cores, 32 threads) and 126 GiB RAM, running Fedora Linux 42. Software versions: R 4.5.2, Python 3.13.11, anndataR 1.1.0, zellkonverter 1.20.1, schard 0.0.1, anndata 0.11.4, rhdf5 2.54.1.

### Supplementary Table 1

Table 1: This table indicates whether a software tool (anndataR, schard and zellkonverter with the native R reader option) provides certain functionality. For each allowed data type for each AnnData slot, we detail whether the software is capable of 1) representing the data type in R, 2) converting it to an appropriate SingleCellExperiment or Seurat object slot and 3) writing the data type back to an H5AD file.
**Legend:**
`x`: the software tool can represent, convert, or write this data type
`/`: the software tool can not represent, convert, or write this data type
`-`: the specific data type is not supported in a SingleCellExperiment or Seurat object.

| | | **anndataR (this paper)** | | | | **schard** | | | | **zellkonverter (native R reader)** | | | |
|---|---|---|---|---|---|---|---|---|---|---|---|---|---|
| **slot** | **datatype** | **in R** | **to SCE** | **to Seurat** | **to H5AD** | **in R** | **to SCE** | **to Seurat** | **to H5AD** | **in R** | **to SCE** | **to Seurat** | **to H5AD** |
| **X** | integer__matrix | x | x | x | x | / | x | x | / | / | x | / | / |
| | integer__csparse | x | x | x | x | / | x | x | / | / | x | / | / |
| | integer__rsparse | x | x | x | x | / | x | x | / | / | x | / | / |
| | float__matrix | x | x | x | x | / | x | x | / | / | x | / | / |
| | float__csparse | x | x | x | x | / | x | x | / | / | x | / | / |
| | float__rsparse | x | x | x | x | / | x | x | / | / | x | / | / |
| | float__matrix__nas | x | x | x | x | / | x | x | / | / | x | / | / |
| | float__csparse__nas | x | x | x | x | / | x | x | / | / | x | / | / |
| | float__rsparse__nas | x | x | x | x | / | x | x | / | / | x | / | / |
| **layers** | integer__matrix | x | x | x | x | / | / | / | / | / | x | / | / |
| | integer__csparse | x | x | x | x | / | / | / | / | / | x | / | / |
| | integer__rsparse | x | x | x | x | / | / | / | / | / | x | / | / |
| | float__matrix | x | x | x | x | / | / | / | / | / | x | / | / |
| | float__csparse | x | x | x | x | / | / | / | / | / | x | / | / |
| | float__rsparse | x | x | x | x | / | / | / | / | / | x | / | / |
| | float__matrix__nas | x | x | x | x | / | / | / | / | / | x | / | / |
| | float__csparse__nas | x | x | x | x | / | / | / | / | / | x | / | / |
| | float__rsparse__nas | x | x | x | x | / | / | / | / | / | x | / | / |
| | float__matrix__3d | x | - | - | x | / | - | - | / | / | - | - | / |
| | integer__matrix__3d | x | - | - | x | / | - | - | / | / | - | - | / |
| **obs & var** | categorical | x | x | x | x | / | / | / | / | / | x | / | / |
| | categorical__ordered | x | x | x | x | / | / | / | / | / | x | / | / |
| | categorical__nas | x | x | x | x | / | / | / | / | / | x | / | / |
| | categorical__ordered__nas | x | x | x | x | / | / | / | / | / | x | / | / |
| | string__array | x | x | x | x | / | x | x | / | / | x | / | / |
| | dense__array | x | x | x | x | / | x | x | / | / | x | / | / |
| | integer__array | x | x | x | x | / | x | x | / | / | x | / | / |

| | | | | | | | | | | | | | |
|---|---|---|---|---|---|---|---|---|---|---|---|---|---|
| | nullable_integer_array | x | x | x | x | / | / | / | / | / | x | / | / |
| | boolean_array | x | x | x | x | / | x | x | / | / | x | / | / |
| | nullable_boolean_array | x | x | x | x | / | / | / | / | / | x | / | / |
| **obsm** | integer_matrix | x | x | x | x | / | x | x | / | / | / | / | / |
| | integer_csparse | x | x | x | x | / | / | / | / | / | / | / | / |
| | integer_rsparse | x | x | x | x | / | / | / | / | / | / | / | / |
| | float_matrix | x | x | x | x | / | x | x | / | / | / | / | / |
| | float_csparse | x | x | x | x | / | / | / | / | / | / | / | / |
| | float_rsparse | x | x | x | x | / | / | / | / | / | / | / | / |
| | float_matrix_nas | x | x | x | x | / | x | x | / | / | / | / | / |
| | float_csparse_nas | x | x | x | x | / | / | / | / | / | / | / | / |
| | float_rsparse_nas | x | x | x | x | / | / | / | / | / | / | / | / |
| | dataframe | x | x | - | / | / | / | - | / | / | / | - | / |
| | float_matrix_3d | x | x | - | x | / | / | - | / | / | / | - | / |
| | integer_matrix_3d | x | x | - | x | / | / | - | / | / | / | - | / |
| **varm** | integer_matrix | x | x | x | x | / | / | / | / | / | x | / | / |
| | integer_csparse | x | x | x | x | / | / | / | / | / | x | / | / |
| | integer_rsparse | x | x | x | x | / | / | / | / | / | x | / | / |
| | float_matrix | x | x | x | x | / | / | / | / | / | x | / | / |
| | float_csparse | x | x | x | x | / | / | / | / | / | x | / | / |
| | float_rsparse | x | x | x | x | / | / | / | / | / | x | / | / |
| | float_matrix_nas | x | x | x | x | / | / | / | / | / | x | / | / |
| | float_csparse_nas | x | x | x | x | / | / | / | / | / | x | / | / |
| | float_rsparse_nas | x | x | x | x | / | / | / | / | / | x | / | / |
| | dataframe | x | - | - | / | / | - | - | / | / | - | - | / |
| | float_matrix_3d | x | x | - | x | / | / | - | / | / | / | - | / |
| | integer_matrix_3d | x | x | - | x | / | / | - | / | / | / | - | / |
| **obsp** | integer_matrix | x | x | x | x | / | / | / | / | / | / | / | / |
| | integer_csparse | x | x | x | x | / | / | / | / | / | x | / | / |
| | integer_rsparse | x | x | x | x | / | / | / | / | / | x | / | / |
| | float_matrix | x | x | x | x | / | / | / | / | / | / | / | / |
| | float_csparse | x | x | x | x | / | / | / | / | / | x | / | / |
| | float_rsparse | x | x | x | x | / | / | / | / | / | x | / | / |
| | float_matrix_nas | x | x | x | x | / | / | / | / | / | / | / | / |
| | float_csparse_nas | x | x | x | x | / | / | / | / | / | x | / | / |
| | float_rsparse_nas | x | x | x | x | / | / | / | / | / | x | / | / |
| | float_matrix_3d | x | x | - | x | / | / | - | / | / | / | - | / |
| | integer_matrix_3d | x | x | - | x | / | / | - | / | / | / | - | / |
| **varp** | integer_matrix | x | x | - | x | / | / | - | / | / | / | - | / |
| | integer_csparse | x | x | - | x | / | / | - | / | / | x | - | / |
| | integer_rsparse | x | x | - | x | / | / | - | / | / | x | - | / |
| | float_matrix | x | x | - | x | / | / | - | / | / | / | - | / |
| | float_csparse | x | x | - | x | / | / | - | / | / | x | - | / |
| | float_rsparse | x | x | - | x | / | / | - | / | / | x | - | / |
| | float_matrix_nas | x | x | - | x | / | / | - | / | / | / | - | / |
| | float_csparse_nas | x | x | - | x | / | / | - | / | / | x | - | / |

| | | | | | | | | | | | | | |
|---|---|---|---|---|---|---|---|---|---|---|---|---|---|
| | float__rsparse__nas | x | x | - | x | / | / | - | / | / | x | - | / |
| | float__matrix__3d | x | x | - | x | / | / | - | / | / | / | - | / |
| | integer__matrix__3d | x | x | - | x | / | / | - | / | / | / | - | / |
| **uns** | categoricals | x | x | x | x | / | / | / | / | / | x | / | / |
| | matrices | x | x | x | x | / | / | / | / | / | x | / | / |
| | scalars | x | x | x | x | / | / | / | / | / | x | / | / |
| | nested values | x | x | x | x | / | / | / | / | / | x | / | / |
| | nan | x | x | x | x | / | / | / | / | / | x | / | / |
| | None | x | x | - | / | / | / | - | / | / | / | - | / |
